# Generative prediction of causal gene sets responsible for complex traits

**DOI:** 10.1101/2025.04.17.649405

**Authors:** Benjamin Kuznets-Speck, Buduka K. Ogonor, Thomas P. Wytock, Adilson E. Motter

**Affiliations:** Department of Physics and Astronomy, Northwestern University, Evanston, IL, 60208; Center for Network Dynamics, Northwestern University, Evanston, IL, 60208; Department of Engineering Sciences and Applied Mathematics, Northwestern University, Evanston, IL, 60208; Northwestern Institute on Complex Systems, Northwestern University, Evanston, IL, 60208; National Institute for Theory and Mathematics in Biology, Evanston, IL, 60208; Chemistry of Life Processes Institute, Northwestern University, Evanston, IL, 60208

## Abstract

The relationship between genotype and phenotype remains an outstanding question for organism-level traits because these traits are generally *complex*. The challenge arises from complex traits being determined by a combination of multiple genes (or loci), which leads to an explosion of possible genotype-phenotype mappings. The primary techniques to resolve these mappings are genome/transcriptome-wide association studies, which are limited by their lack of causal inference and statistical power. Here, we develop an approach that leverages transcriptional data endowed with causal information and a generative machine learning model to strengthen statistical power. Our implementation of the approach—dubbed TWAVE—includes a variational autoencoder trained on human transcriptional data, which is incorporated into an optimization framework. Given a trait phenotype, TWAVE generates expression profiles, which we dimensionally reduce by identifying independently varying generalized pathways (eigengenes). We then conduct constrained optimization to find causal gene sets that are the gene perturbations whose measured transcriptomic responses best explain trait phenotype differences. By considering several complex traits, we show that the approach identifies causal genes that cannot be detected by the primary existing techniques. Moreover, the approach identifies complex diseases caused by distinct sets of genes, meaning that the disease is polygenic *and* exhibits distinct subtypes driven by different genotype-phenotype mappings. We suggest that the approach will enable the design of tailored experiments to identify multi-genic targets to address complex diseases.

**Significance summary:** Researchers have long sought to bridge the gap between phenotypes and the genotypes that cause them. This gap remains open because current methods focus on associating phenotypes to a combinatorially explosive number of genotypic possibilities, resulting in a loss of statistical power. We overcome this limitation by employing transcriptomic data from complex, polygenic, human diseases combined with measured transcriptomic responses to gene perturbations in cell lines. The former data allow us to perform generative modeling and dimensional reduction to map transcriptome to phenotype, while the latter incorporate causal information regarding how gene regulation shapes phenotype. We predict sets of genes that explain the emergence of complex traits, which suggest possible multi-target disease treatments.

## Introduction

Complex traits are polygenic, orchestrated by networks of correlated genes that work together to produce phenotypic variation [1, 2, 3]. An outstanding question in the study of such traits is the identification of the specific combinations of gene variants that give rise to the different phenotypic expressions [4, 5, 6, 7, 8, 9, 10]. Association studies have been performed to search for genetic loci significant to a complex trait phenotype by conducting hypothesis tests on *individual* genetic loci, from which *independent* mutations/genes are statistically associated with the phenotype in question [11, 12, 13, 14, 15]. We innovate on these techniques by developing a framework to *jointly* predict *sets* of genes while accounting for collective behavior not captured by statistical tests on individual genes.

Association studies, such as genome/transcriptome-wide association studies (GWAS/TWAS), have been broadly adopted in over 5,700 studies and 3,300 traits as of 2021 [12]. A common critique of these studies is that they have low statistical power due to the combinatorial explosion in the number of gene sets that must be tested [11, 13, 16]. Post-GWAS/TWAS analyses such as *fine-mapping* attempt to address this limitation by considering the correlation structure of the genetic data [17, 18, 19, 20, 21]. However, they rely on an initial association study to select what variants to fine-map, potentially leaving behind genes that would be significant collectively but have low independent effect size. In the framework presented here, we develop and apply an approach that considers all genes simultaneously, regardless of individual effect size. The framework combines generative machine learning, dimensionality reduction, and constrained optimization (Fig. 1).

**Figure 1:**
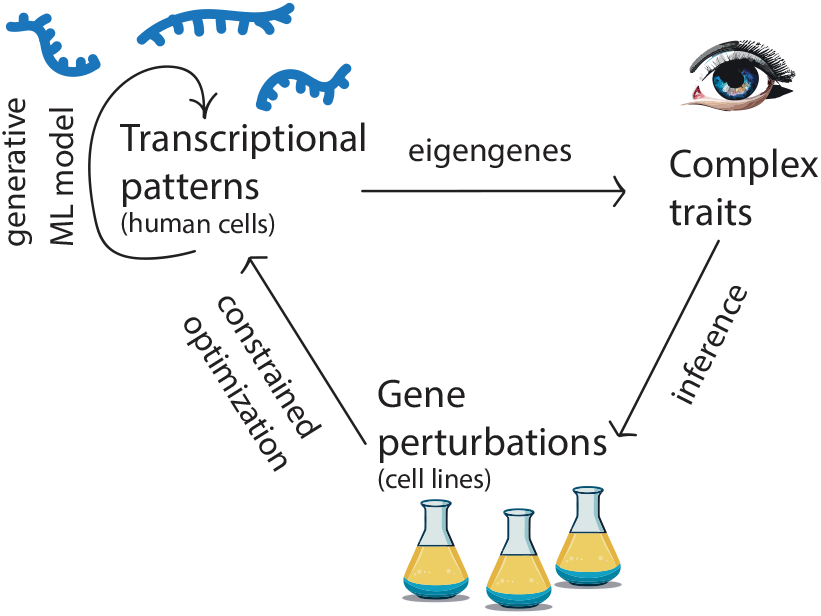
Schematic of the proposed approach. Synthetic transcriptomic profiles are generated from a machine learning model that learns from RNA-Seq data on complex traits. The generated data are projected onto eigengenes, which are linear combinations of genes that vary independently and retain important gene correlations that differentiate between complex trait phenotypes. From here, gene perturbations whose transcriptional responses bridge the gap between trait phenotypes are found by constrained optimization.

A key aspect of our approach is the use of increasingly available trait-labeled transcriptomic data from bulk and single-cell RNA-Seq experiments, which contends with the biological networks that influence complex traits [22, 23]. To better extract patterns from our transcriptional data, we develop the Transcriptome-Wide conditional Variational auto-Encoder (TWAVE), a machine learning model that generates denoised transcriptional profiles for the relevant phenotypes. To understand how regulatory changes affect expression levels, we also incorporate complementary data on transcriptional response to gene perturbations (knockdowns and overexpressions) [24]. These transcriptional data are dimensionally reduced while maintaining causal information, which we achieve using the concept of eigengenes, to facilitate the optimization over gene perturbations [25]. Together, these data sources allow us to explore how the regulatory network drives phenotypic changes without prior knowledge of network structure.

We focus on the human disease traits in Table 1. Throughout, we take care to distinguish between *traits* (e.g., eye color) and their *phenotype* variants (e.g., blue, brown, green), and we consider traits that have a baseline and variant phenotype. Moreover, we interpret the states associated with each variant as defined by distinct attractors of the gene regulatory network [26, 27, 28]. The problem of identifying the genes that cause a trait phenotype can thus be mapped to an optimization over combinations of transcriptional perturbations that steer transcriptomic states from baseline to variant attractors and vice-versa. The resulting framework reveals groups of gene perturbations that most influence phenotypic variation, pinpointing the molecular underpinnings that determine complex traits.

**Table 1:** Gene Expression Omnibus and DepMap datasets. The columns represent (from left to right) the traits considered, the tissues of origin, the type of sequencing, the number of samples of the baseline and variant phenotypes, and the GEO or DepMap series accession numbers.

| Complex trait | Tissue | Seq. Type | $N_{\text{baseline}}$ | $N_{\text{variant}}$ | GEO Series |
| --- | --- | --- | --- | --- | --- |
| Allergic asthma | PBMCs | bulk | 277 | 166 | GSE96783 |
| IBD | gastrointestinal tissue | bulk | 461 | 2029 | GSE193677 |
| Food Allergy | CD4+ T-cells | bulk | 71 | 63 | GSE114065 |
| Cancer metastasis | metastasis: breast → lung | single | 1170 | 1274 | GSE202695 |
| AMD | macular retina & RPE | bulk & single | 433 | 104 | GSE135092 |
| Type 1 diabetes | CD4+ T-cells | single | 557 | 2502 | GSE182870 |
| NSCLC | blood platelets | bulk | 376 | 400 | GSE89843 |
| Simple trait | Tissue | Seq. Type | $N_{\text{baseline}}$ | $N_{\text{variant}}$ | GEO Series |
| MODY3 | differentiated ESCs | single | 113 | 158 | GSE129653 |
| Complex trait | Tissues | Seq. Type | $N_{\text{baseline}}$ | $N_{\text{variant}}$ | DepMap Series |
| Metastasis | 86 cancers, 34 lineages | single | 838 | 447 | 24Q2 |

## Results

### Generating complex trait transcriptomes

Identifying *sets* of differentially expressed genes is complicated by the fact that transcriptional measurements include a large number of genes and a comparatively small number of samples. It is precisely this feature that makes it challenging to distinguish real biological differences from random variance when using statistical tests that treat genes as independent variables. Our method recognizes that genes operate in concert rather than independently to orchestrate cell function, which transforms the problem into learning an effective representation of these relationships from data. Fig. 2 presents TWAVE, which solves this problem by looking at the data as a whole, using a neural network encoder to embed high-dimensional gene expression profiles onto a low-dimensional latent space (*Z*), where data points can be classified and new representative points can be generated. Points in the latent space, including newly generated ones, are decoded back up to the full gene expression space as in Fig. 2A. The model (consisting of the encoder, decoder, and latent space classifier) is trained with a combination of three loss terms. The first term accounts for how accurately the autoencoder can reconstruct the original data. The second term is the Kullback-Leibler (KL) divergence loss, which regularizes *Z* so that each dimension contributes roughly equally to the overall variance in the latent space. The third term is a classification loss to mold the structure of the latent space so that different phenotypes of the trait (baseline and variant for example) can be distinguished by their transcriptional states, as detailed in Materials and Methods.

**Figure 2:**
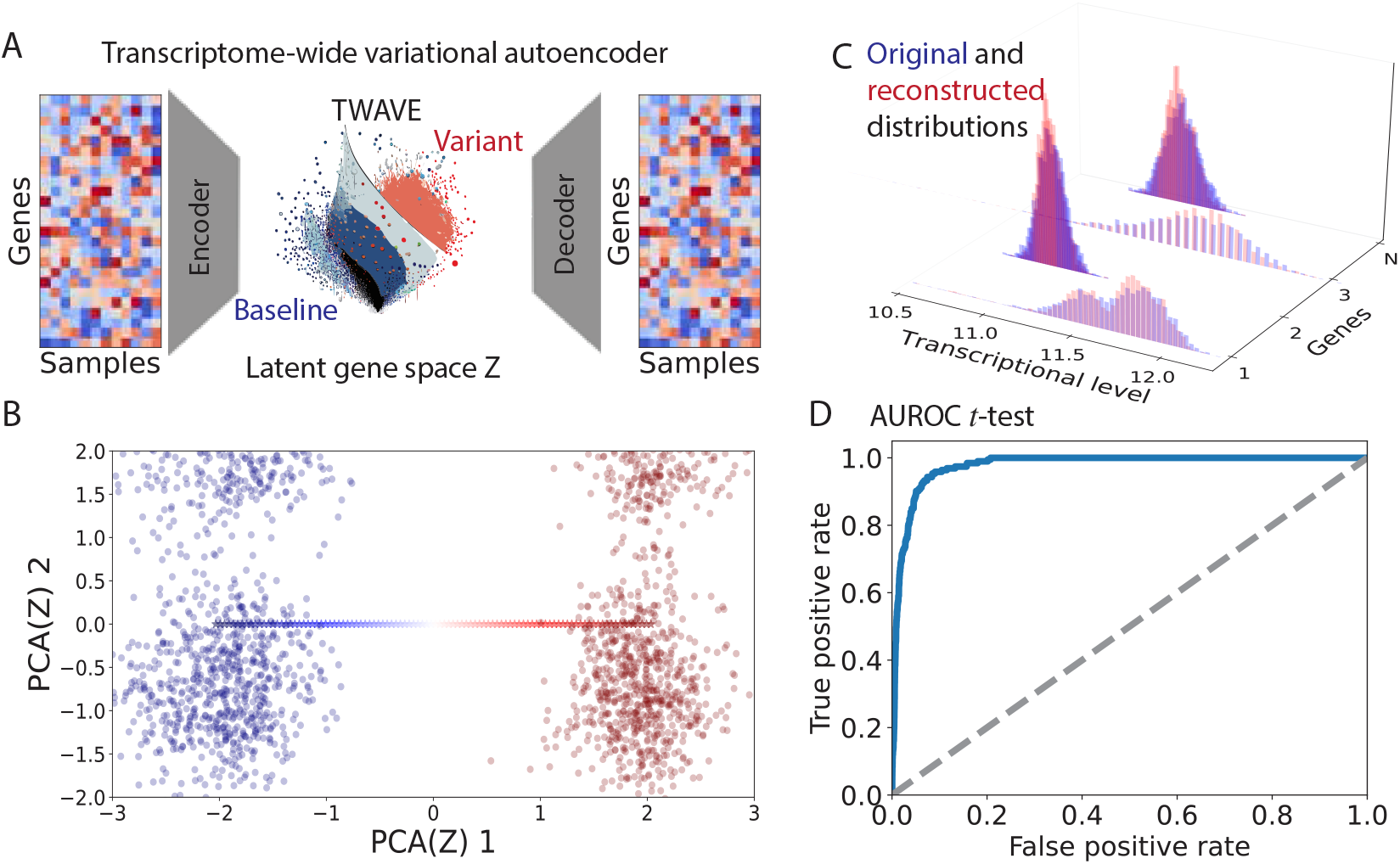
TWAVE construction and validation, presented for the inflammatory bowel disease trait. (*A*) TWAVE architecture, where gene expression profiles are projected onto a low-dimensional latent space (*Z*) and subsequently reconstructed with a neural network decoder. (*B* ) First two principal components (PC) of the latent space *Z*, showing a clear separation of complex trait phenotypes (baseline and variant). A linear interpolation between the means of the two populations in the latent space (blue-to-red stars) falls along the first principal component. (*C* ) Comparison between original (blue) and TWAVE-reconstructed (red) distributions of gene expression for 4 different genes, conveying strong agreement. (*D* ) Receiver operating characteristic for significant gene associations in the reconstructed versus original data, confirming that TWAVE retains significant associations found in the original data.

Fig. 2B shows that transcriptional measurement associated with the baseline and variant populations (blue and red clusters) segregate to coordinates along the first principal component, which is a natural outcome of the latent space learned by TWAVE. Here, we use the representative case of inflammatory bowel disease to illustrate the construction of TWAVE, but we observe similar performance among the other traits we considered in Table 1. The figure also shows that a linear interpolation between the two clusters in *Z*-space lies along the first principal component, which accounts for the largest fraction of variation in the data. Fig. 2C demonstrates a close agreement between the original gene expression distributions and the reconstructions from TWAVE. To test whether our auto-encoder retains associations between genes and the complex trait of interest, we compare the differentially expressed genes identified by *t*-tests on both the original and reconstructed expression profiles using the area under the receiver operating characteristic curve (AUROC). Fig. 2D shows this curve, which is constructed by: 1) arranging both sets of genes from smallest to largest in terms of their *p*-values, 2) varying the threshold at which genes are statistically significant, and 3) counting the fraction of significant genes in the original data that are selected by TWAVE (true positive rate) as a function of the fraction of genes selected by TWAVE that are *not* significant in the original data (false positive rate). For all traits, we find that the AUROC approaches 1, indicating that the sets of differentially expressed genes identified by TWAVE and within the original data are nearly identical. Full technical details concerning the construction of TWAVE are provided in Materials and Methods.

For each of the complex trait datasets described in Table 1, we use TWAVE to estimate the distributions of the transcriptional data in the latent space arising from the baseline and variant phenotypes, while retaining the fundamental features that distinguish the two populations. We then draw points from these distributions in the latent space, decoding them as depicted in Fig. 2A. By choosing an equal number of each phenotype (baseline and variant), estimates of the distributions from the data are equally precise for each of these trait phenotypes. It is instructive to compare against the method of “extreme pseudo-sampling,” in which the latent space of a variational auto-encoder (VAE) is sampled randomly [29, 30]. Our method is different from extreme pseudo-sampling in two crucial ways. First, TWAVE employs a conditional VAE (i.e., it includes a latent space explicitly trained to classify between baseline and variant). Second, TWAVE draws from a probability distribution in the latent space associated with the trait phenotype label instead of drawing randomly from *any* state in the latent space. Overall, TWAVE allows us to make the most from limited transcriptional data, drawing new representative samples in a way that would be unfeasible without generative modeling.

### Causal dimensions of complex trait variation

We select the *eigengenes* that are most determinative of the trait phenotypes (i.e., are causal). Conceptually, eigengenes are weighted combinations of genes that vary in concert within the eigengene, but any given eigengene can vary independently of the others. Mathematically, they are eigenvectors of the TWAVE-generated gene expression matrix *Y* and form an orthogonal basis ***e***_*i*_ in gene expression space [25]. This basis corresponds to the columns of the unitary matrix *U* = [***e***_0_, …, ***e***_*i*_, …, ***e***_*n*_] in the singular value decomposition *Y* ^*T*^ = *U* Σ*V* ^*T*^ , where Σ is a diagonal matrix of singular values in descending order and *V* ^*T*^ contains the left eigenvectors of *Y* ^*T*^ . For each dataset, we perform singular value decompositions of the *m* × *n* matrix *Y* , where *m* is the number of sample expression profiles in the TWAVE dataset and *n* the number of genes in each sample.

We proceed to determine which eigengenes are most likely to capture differences between the baseline and variant trait phenotype by adapting Bayesian fine-mapping [17] to eigengenes. This procedure seeks a small set of *r* eigengenes that can accurately distinguish between the phenotypes according to the posterior inclusion probability, which quantifies how well a proposed set of eigengenes explains the data (i.e., how causal the set is). The fine-mapping procedure starts by projecting the data onto the *d* = 200 eigengenes with the largest singular values *X* = *Y U*_*n*×*d*_. This choice of *d* ensures that the set of eigengenes from which the causal set is selected accounts for the large majority of the variance, as shown in Fig. S1. A logistic regression model over eigengenes is then fit using the expression profiles and associated trait phenotype labels, achieving high accuracy, F1-score, and recall, for all datasets analyzed. We also show that a maximum likelihood estimator for the posterior distribution of causal eigengenes can be formed from: 1) the odds ratios *ζ* = log[*ρ/*(1 − *ρ*)] from the logistic regression, where *ρ* is the probability of a data point belonging to the variant class; and 2) the projected expression matrix *X*. To reduce from *d* to *r* eigengenes, we perform Markov chain Monte Carlo (MCMC) sampling to maximize the likelihood of causal eigengenes given the regression data. For each of these *d* eigengenes, we evaluate the posterior inclusion probability that each eigengene is causal *p*(***e***_*i*_ causal | data = {*X, ζ*}) and build sets from the top *r* = 50 causal eigengenes (*r* = 10 for allergic asthma) with the largest posterior inclusion probability. We show *p*(***e***_*i*_|*X, ζ*) for the first *d* principal components of *Y* in Fig. 3A, ordered from non-causal (*p* = 0) to causal (*p* = 1).

**Figure 3:**
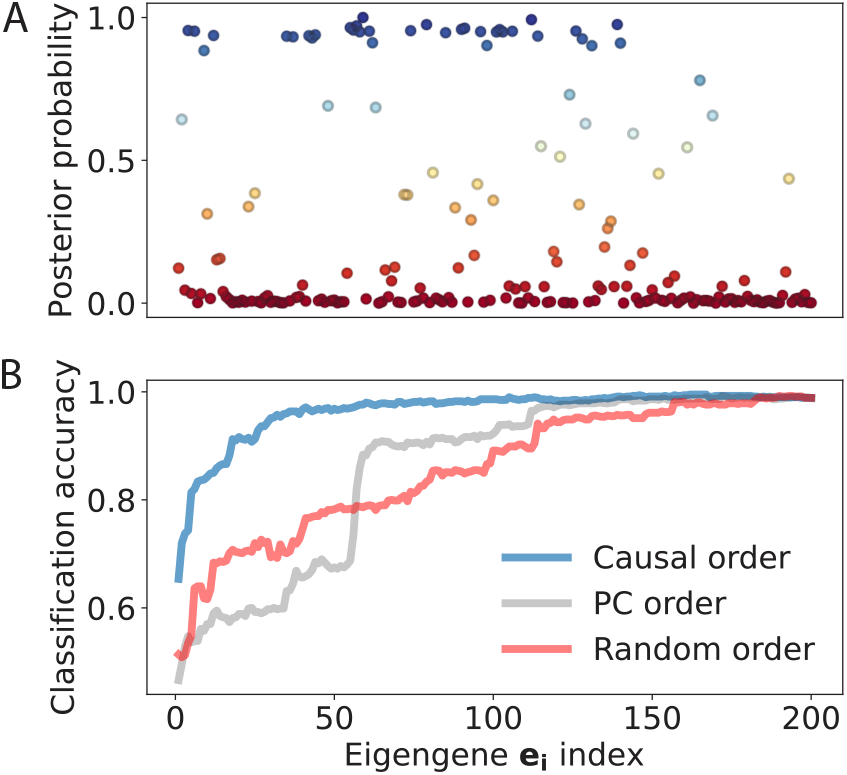
Dimension reduction by selecting the most causal eigengenes for the inflammatory bowel disease trait. (*A*) Posterior inclusion probability for eigengenes to be causal arranged in descending order of their singular values. (*B* ) Classification accuracy from logistic regression on data projected onto an increasing subset of the top *d* eigengenes arranged in descending order of the posterior probability of being causal (blue), in descending order of the singular values (gray), and randomly (red).

As a validation test for the selected eigengenes, we perform the logistic regressions on *X* summarized in Fig. 3B. The regression accuracy for including data projected onto the first *i* eigengenes sorted in order of causality *p*(***e***_*i*_|*X, ζ*) quickly climbs to above 0.9 within *r* top eigenvectors. This validates our choice of keeping *r* eigengenes in our dimensionality reduction. On the other hand, including eigengenes in principal component order (i.e., ordered by the fraction of variance that aligns along each eigenvector), yields significantly poorer classification results. Arranging the eigengenes randomly can actually produce a better result than arranging them in order of the singular values when keeping less than 60 eigengenes, but principal component ordering outperforms the random one as the dimension of this reduced space is increased.

### Complex trait transitions via eigengene perturbations

#### Constrained optimization

To implicate genes responsible for transitioning between the baseline and variant phenotypes, we explore extensive data on transcriptional responses to gene perturbations. Specifically, we define a perturbation response matrix, *B*, whose rows represent eigengenes in the original dataset and whose columns are average transcriptional responses to a transcriptional perturbation (one response profile for each column). This matrix consists of 10% overexpressions and 90% knockdowns (most of the latter are implemented through RNA interference), as indicated in Dataset S1. With this matrix in hand, we investigate which combinations of perturbations can cause the baseline transcriptional profile to match the variant, and vice-versa. Formally, this question is answered by solving the following constrained optimization problem:

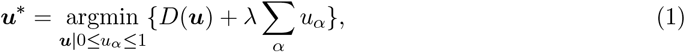

where

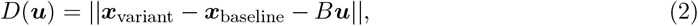

and || · || denotes the Euclidean distance. The choice of Euclidean distance reflects our assumptions 1) that there is a single phenotype for each transcriptional state and 2) that differences in the expression of each eigengene are equally likely to contribute to phenotypic differences. We discuss alternative choices of the distance metric in the Supplementary Information (SI). Here, perturbations add with weight ***u*** in causal eigengene space to transition from the state ***x***_baseline_ to the state ***x***_variant_ (Fig. 4A). Before considering transitions between individual states in the baseline and variant clusters, we consider transitions between *average* baseline and variant states.

**Figure 4:**
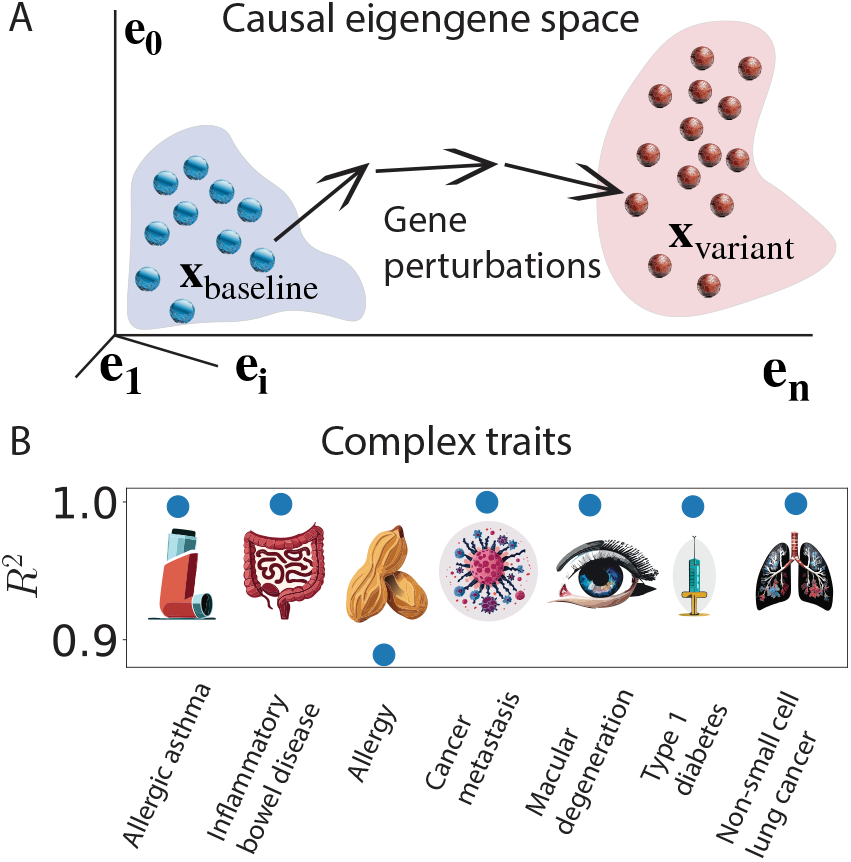
Attributing complex trait phenotypes to sets of genes. (*A*) Identification of gene sets that differ between baseline and variant trait phenotypes, where blue and red dots represent individual states. A state in the baseline cluster (blue background) transitions to a state in the variant cluster (red background) upon targeted transcriptional perturbations. (*B* ) Coefficient of determination for controlled transitions between phenotypes of complex traits, where *R*^2^ close to 1 indicates that the final transcriptional state approaches that of the target phenotype.

Since we expect the baseline and variant phenotypes to be stable with respect to the fluctuations inherent to transcription, the closer we approach states known to belong to a given phenotype transcriptionally, the more likely that state is to exhibit that phenotype. We quantify this likelihood with the coefficient of determination,

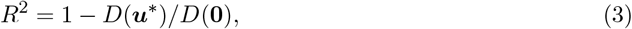

where *R*^2^ ≤ 1 and *R*^2^ close to 1 indicates high efficacy. In particular, *R*^2^ *>* 0.5 indicates that the distance between the two states has been at least halved. The relationship between distance and cell behavior becomes less precise in the full high-dimensional expression space because there are many points that are a given distance away from a target point. Thus, reducing the number of dimensions of our problem by working with a select set of eigengenes is crucial, which is implemented through our choice of matrix *B*. We use the Python function minimize from SciPy (which implements the L-BFGS-B method) to solve the constrained optimization problem in Eq. 1 for all of our RNA-Seq datasets. The optimal ***u***^*^ yields *R*^2^ ≈ 1 for all complex traits we consider, namely allergic asthma, inflammatory bowel disease, food allergy, cancer metastasis (where baseline and variant refer to primary and metastatic tumors), age related macular degeneration, type-1 diabetes, and non-small cell lung cancer (Fig. 4B). This is further confirmed by examining the selected perturbations for individual trait phenotypes and noting that they generally point towards the phenotype expression in question.

Table 2 shows the genes with the top 12 perturbation weights in ***u***^*^ for allergic asthma along with a brief annotation of their function. Many of these top-selected genes have been implicated in allergic asthma, lung and airway function, and inflammation and immunity. Mutations in *BMPR2*, for example, have been shown to cause asthma-like symptoms and pulmonary hypertension in response to mild antigens in the airway [34, 35]. In addition, *TCF7* promotes T-cell differentiation to Th2 or memory T-cells [36], consistent with allergic asthma being the result of immune system dysregulation. Other identified genes, such as *TARDBP, TENT4B*, and *HNRNPL*, have not been previously implicated in allergic asthma. However, these genes are associated with RNA metabolism and modifications including alternative splicing and poly-A tail alteration, which in turn are related to immune response [46]. In particular, *TARDBP* (*TDP-43* ) has been shown to regulate alternative splicing and alternative polyadenylation in CD8+ T-cells, and specific RNA splicing and polyadenylation events depend on the presence of *TARDBP* during CD8+ T-cell co-stimulation [31]. *TENT4B* is involved in mRNA stabilization, influencing B-cell proliferation and the cellular response to viral infections [32], whereas *HNRNPL* participates in the regulation of inflammatory responses, particularly through its interaction with long non-coding RNAs and its role in regulating *TNF-α* transcription [44, 45]. We performed the optimization between the average baseline and variant states for all 6 other complex traits shown in Fig. 4B, and refer the reader to Supplementary Tables S1-S6 for the top selected genes for each trait.

**Table 2:**
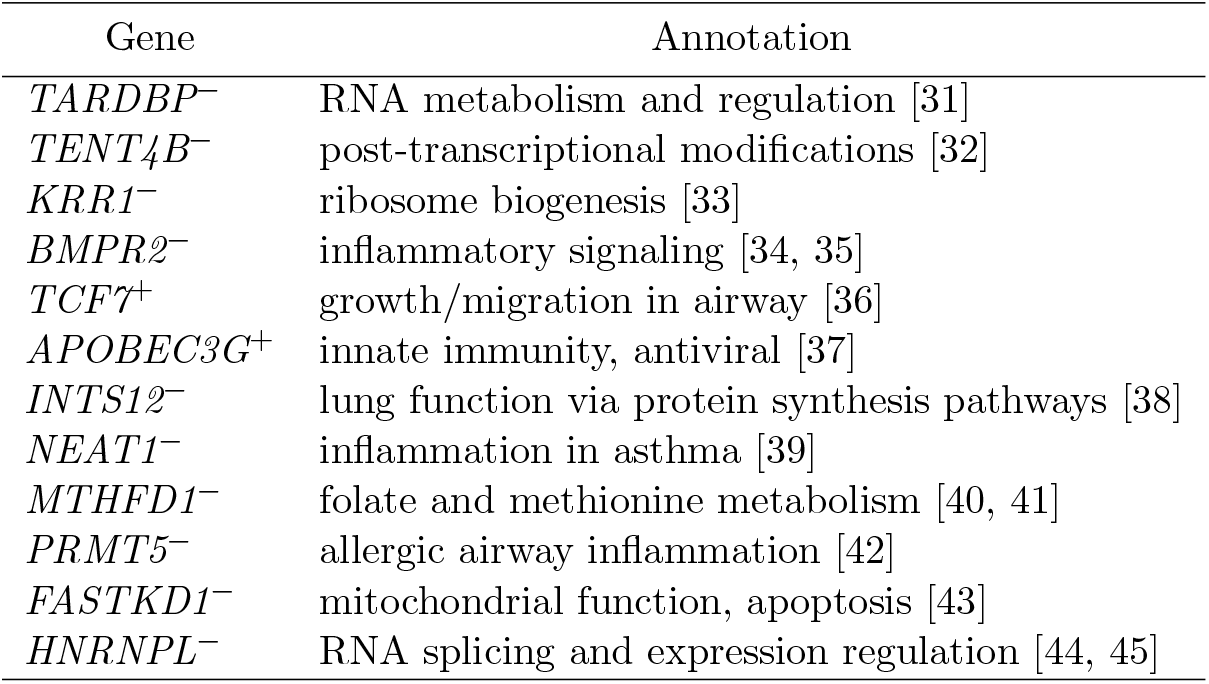
Allergic asthma transcriptional perturbations (GSE96783)

#### Optimization for individual baseline-variant states

We perform optimizations across individual baseline and variant pairs to account for the fact that the measured transcriptional signatures of a given a phenotype vary heterogeneously across cell samples and individuals (Fig. 4A). An analogous approach in TWAS would require sub-sampling an already small number of measurements, leading to a large uncertainty in the variance and an inability to detect baseline-variant differences [47]. By recasting the hypothesis test as an optimization problem, we avoid this issue using information on how the regulatory network responds to perturbations. This reformulation allows us to investigate how the gene perturbations responsible for a trait may change across individual baseline-variant pairs. We break down our strategy into two steps: 1) find the the optimal set of perturbations for a large number of baseline-variant state pairs and 2) compare the observed co-occurrence of perturbation pairs with a null model. The null model is designed to preserve both the frequency with which each perturbation is selected and the number of perturbations needed for each pair of states.

The transition between any ***x***_*i*_ and ***x***_*j*_ can be induced by applying the perturbation

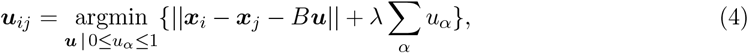

where the variant state is ***x***_*i*_ and the baseline state is ***x***_*j*_. We solve this optimization problem over a range of different *λ* values, taking the largest *λ* (the sparsest solution) such that *R*^2^ = 1 − *D*(***u***_*ij*_)*/D*(**0**) *>* 0.99. This is repeated for *N* = 2500 randomly selected pairs in the forward (baseline-to-variant) and reverse directions. We then construct a bipartite network represented by the (adjacency) matrix

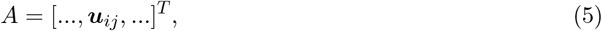

where the columns of *A* are perturbed genes and the rows are different accepted *i, j* pairs. We take the dot-product between the columns *µ* and *ν* of *A* to get the frequency *f*_*µν*_ at which the corresponding perturbed genes co-occur in the same ***u***_*ij*_:

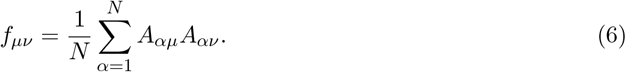

To identify statistically significant pairs *µ, ν*, we compare *f*_*µν*_ to the frequency at which *µ* and *ν* occur together in a null model of a random maximum entropy graph whose row and column sums are fixed to that of *A* [48, 49]. The expected co-occurrence frequency in the null model is

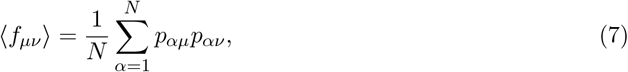

where *p*_*ij*_ is the probability of an edge existing between nodes *i* and *j* of the maximum entropy graph (see Materials and Methods). In addition to ⟨*f*_*µν*_⟩, the standard error *σ*_*µν*_ can be approximated via error propagation using the fact that the probability of edge occurrences are Bernoulli random variables. From these ensemble statistics a z-value can be constructed as *z*_*µν*_ = (*f*_*µν*_ − ⟨*f*_*µν*_⟩)*/σ*_*µν*_.

Now that we have established a null model for the co-occurrence of perturbations in causing/reversing the variant behavior, we can examine the network formed by the statistically significant co-occurrences that deviate from the maximum entropy model. To determine where co-occurrences begin to deviate from the null model, we identify the set of significant pairs with high *z*_*µν*_ above a threshold defined by inspection of the quantile-quantile plot (see SI Fig. S2).

#### Application to allergic asthma

In the case of allergic asthma, pairs with *z*_*µν*_ *>* 20 were kept for analysis. The corresponding network is depicted in Fig. 5A for the baseline-to-variant transition (i.e., the onset of asthma). Each node represents a perturbed gene within a significantly co-occurring pair and is color-coded according to the nature of the perturbation (knockdown or overexpression). Edges represent significant co-occurrences and are color-coded by the expression correlations across responses between the genes they connect. We find that genes with many connections in this network representation, such as *ADAR, PAN3*, and *MAPK1* have been implicated in allergic asthma before [50, 51, 52]. We also optimized over gene perturbations in the reverse direction, as shown in Fig. 5B. Among the genes featured in this network, we again find several linked to allergic inflammation: *SUZ12* inhibition is associated with the reduction of allergic inflammation through is role in the protein complex PRC2, *JAK2* inhibitors have been proposed to alleviate asthma because of *JAK2* ‘s role in the JAK-STAT signaling pathway, and knockdown of *MYC* has been shown to repress ILC2 (immune cell) activity, which in turn reduced airway inflammation and immune hyperresponsiveness [53, 54, 55].

**Figure 5:**
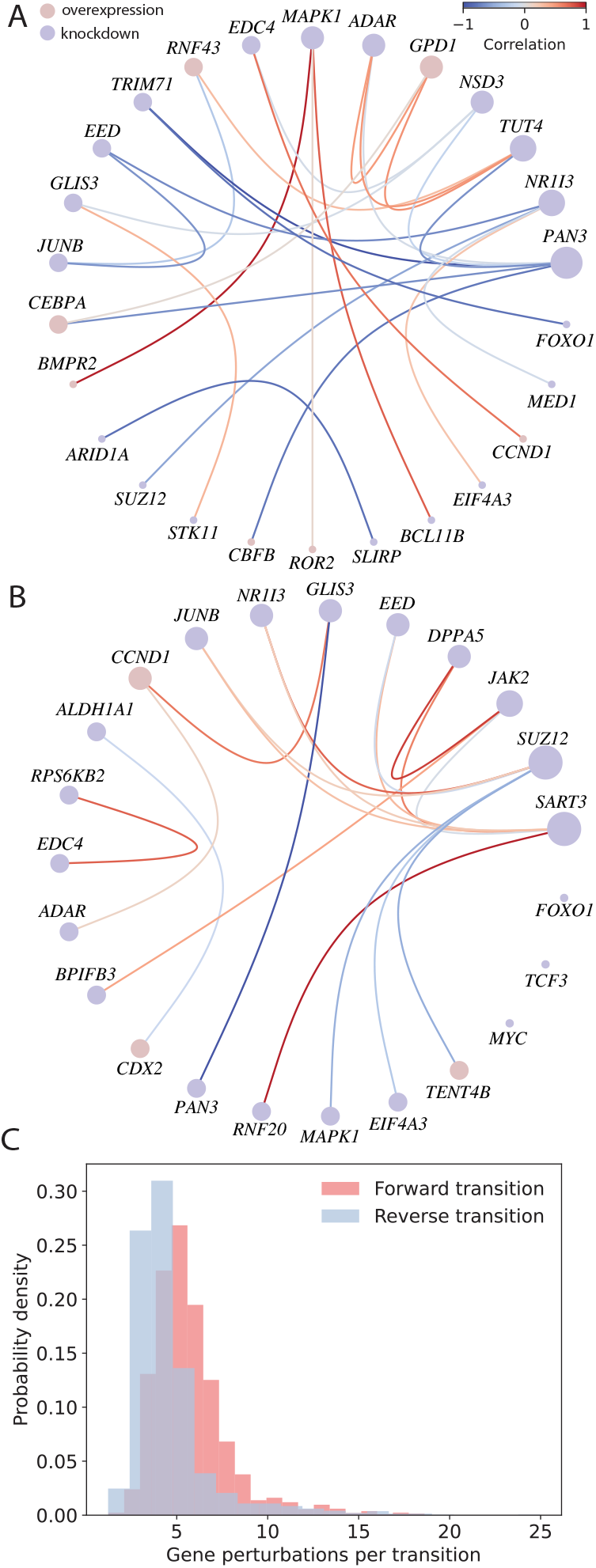
Genes perturbed in transitions between baseline and variant clusters for allergic asthma. *(A, B* ) Gene perturbation co-occurrence networks for forward (A) and reverse (B) transitions. Edges appear between a pair of perturbed genes if they frequently co-occur in successful transitions (i.e., they both occur with high frequency in ***u***_*ij*_ compared to a maximum entropy graph null model). The edges are colored by gene-gene correlation in the perturbation response dataset. The nodes are sized proportionally to the number of edges and color-coded according to whether the perturbation is a knockdown (blue) or overexpression (red). (*C* ) Histograms of the number of gene perturbations required to induce the forward transition (red) and the reverse transition (blue).

Note that many of the genes in the forward and reverse co-occurrence graphs in Fig. 5A, B are *distinct*. In a dynamical network, reversing a perturbation does not generally restore the state of the system, a phenomenon that is accounted for by the bounds placed on ***u*** in Eqs. (1) and (4). These bounds may prevent the same genes from being selected in the forward and reverse directions, which within our approximation reflects the fact that the responses to knockdowns and overexpressions are not exactly anti-aligned. In our case, this phenomenon gives rise to the observation that the genes causing a given trait phenotype are not necessarily the ones that mitigate it. Moreover, as shown in Fig. 5C, the number of perturbations required to make the transition in the forward and reverse directions are also different. Remarkably, it takes a combination of fewer single-gene perturbations to induce a transition in the reverse direction (i.e., from the variant to the baseline state) than in the forward one. This may be because it takes more perturbations to go to a *particular* variant state than to a generic baseline state (“all roads lead to Rome” but not necessarily the inverse). A second possibility is that there is an overrepresentation of certain genes in the library of gene perturbations (see Materials and Methods).

Importantly, both genes involved in the forward direction *and* genes involved in the reverse direction can be associated with the trait because they can promote or reverse a change from the baseline to variant phenotype. For example, the involvement of *MYC* in the reverse direction may be due to its known function in promoting plasticity between transcriptional states and amplifying gene expression overall when overexpressed [56, 57]. Moreover, pluripotent stem cells with *MYC* knocked-down have been shown to decrease allergic reactions in mice by inhibiting T-helper cell immune reaction [58]. Aberrant translation of the *CEBPA* gene, which is implicated in the forward direction, has also been associated with causing bronchial smooth muscle cells—a tissue that plays a key role in asthma—to proliferate faster [59]. Notably, there are genes featured in both the forward and reverse co-occurrence networks that play roles in both promoting and attenuating allergic asthma. For example, *FOXO1* overexpression in mice has been shown to promote allergic asthma through macrophage polarization, Th9 (T-helper 9 cell) differentiation, and regulation of *IRF4* expression, though inhibition of *FOXO1* led to attenuation of immune response and asthmatic inflammation through regulation of *IRF4* [60, 61]. Likewise, the role of *JUNB* depends on the state of other transcription factors. Though *JUNB* significantly influences Th2 (T-helper 2 cell) differentiation and the production of Th2 cytokines, promoting allergic inflammation, it also plays a role in maintaining homeostasis. Specifically, knockdown of *JUNB* limits excessive inflammation by modulating regulatory T-cell differentiation [62].

We also find that, except for *BMPR2* and *TENT4B*, the genes appearing in the co-occurrence graphs from the optimization between *individual* baseline-variant states are distinct from those identified by the *average* variant-baseline optimization (Table 2). The apparent contrast between optimizing over average transcriptional states and individual pairs highlights the fact that there can be multiple paths (defined by different perturbation sets) through which the disease progresses and is mediated and that these paths are not necessarily the ones connecting the average transcriptional states. Consequently, the distinct co-occurring genes in Fig. 5 could potentially relate to different mechanisms playing a role in allergic asthma. We emphasize that such a co-occurrence structure *cannot* be inferred from studying population averages alone, as typically done in GWAS/TWAS [47].

It is instructive to consider the transcription factors that regulate the genes in our perturbation response library (*upstream* transcription factors). Not all upstream transcription factors have transcriptional responses measured in our library, so we use the Enrichr gene set enrichment database to find them. We focus on the case where the upstream factors simultaneously regulate both genes in a co-occurring pair so that a single transcription factor could explain their combined influence on the trait phenotype. For instance, *GATA2, TET2*, and *TWIST1* are enriched for more than one gene co-occurrence pair and are known to influence allergic asthma [63, 64, 65]. The most parsimonious explanation for the enrichment of these transcription factors is that they exert their influence on the phenotype (at least in part) through the genes that appear in our perturbation response dataset. The enrichment of these additional factors shows that we may be able to infer trait-associated genes outside our dataset.

#### Learning across different contexts with TWAVE

Thus far, we have considered 7 complex traits, each based on data from a unique tissue type and previously examined by differential expression. Next, we generalize to contexts where 1) the data (and trait phenotype) in question are associated with multiple disparate tissues and 2) the trait phenotype is caused by a mutation that affects the function of the protein instead of its transcriptional expression.

We consider a phenotype that manifests itself through multiple tissues by studying the trait of cancer metastasis in the cancer dependency map (DepMap) dataset [66]. Because these *pan-cancer* data come from many different cell types, there are confounding variables that render a simple differential expression analysis unable to detect any differentially expressed genes common to the process of metastasis across all tissues. Indeed, we found no statistically significant associations by performing such an analysis. However, using TWAVE, we are able to disentangle the effects of the confounding variables associated with different disease contexts (e.g., cell type, tissue origin, tumor location, and systematic effects) to uncover the common biological mechanisms driving cancer metastasis (see SI Fig. S3). In this case, we again find that our co-occurring perturbation networks contain many genes previously found to promote or mitigate cancer metastasis, including *NF1* knockdown, *SOX5* overexpression, *CBFB* overexpression, *TOX4* knockdown, *PROX1* over-expression, and *EHF* knockdown [67, 68, 69, 70, 71, 72]. An Enrichr search for the co-occurring genes reveals out-of-sample upstream transcription factors that are known to affect metastasis as well, such as *STAT3* and *CTCF* [73, 74]. Although both of these genes appear to be essential for growth [75, 76], which limits opportunities to perturb them in cell-line experiments, one can detect their influence on trait phenotype variation through the pairs of genes in the co-occurrence graph that they regulate.

To examine the scenario in which a causal mutation affects a gene’s protein function but not its transcriptional expression, we consider maturity-onset diabetes of the young type 3 (MODY3). MODY3 is known to be a largely *monogenic* trait caused by mutations to the transcription factor *HNF1A* that impact beta cell function and diabetes in general. Since *HNF1A* is one of the genes perturbed in our perturbation response matrix *B*, MODY3 provides an excellent example where the solution is known. Documented mutations of *HNF1A* alter its protein function, which in turn alters the expression of *other* genes as opposed to its own. In fact, *HNF1A* overexpression appears in 30.2% of baseline-variant pair optimizations. This is compared to the top overall perturbation, *NEAT1* knockdown, appearing 57.4% of the time. However, *HNF1A*, as opposed to *NEAT1*, also appears in the forward co-occurrence network (see SI Fig. S4). Of the top 13 gene perturbations, three of them—*MED1* knockdown, *HNF1A* overexpression, and *GATA2* overexpression—were also in the forward co-occurrence network and have been implicated in diabetic function.

This narrows down the large list of possible genes to a number that could be tested in low-throughput lab experiments. For instance, *MED1* knockout mice show a heightened sensitivity to insulin and an improved glucose tolerance [77]. All of the other genes in the co-occurrence network exhibit transcriptional responses that are highly positively correlated with that of *HNF1A*, meaning that their corresponding column vectors in *B* all point in the same direction. This pattern is markedly different from those observed in the complex traits above, in which we also find transcriptional responses that are negatively correlated and uncorrelated. Among the perturbations correlated with *HNF1A* overexpression is *ALOX5* overexpression, which also impacts beta cell function in diabetes via increased insulin resistance [78, 79].

Finally, we directly compare the genes identified by our method, differential expression, and TWAS in the case of inflammatory bowel disease in SI Fig. S5. We find that the only 8% of the differentially expressed genes identified in the dataset [80] overlap with the TWAS genes, which is reflective of the challenges inherent to reconciling results produced by different approaches. Applying our method, we find that 36% of the genes participating in over 54% of the solutions to Eq. (4) overlap with TWAS. We emphasize that this improvement by our method occurs because it accounts for the downstream impacts of the gene perturbations through the *B* matrix, which naturally filters out spurious differentially expressed genes.

## Discussion

The approach presented here leverages existing transcriptomic data to address the challenges that complex traits pose to traditional mutation-association screening methods [13]. We implement this by identifying generalized cellular pathways (eigengenes) relevant to a complex trait and by calculating optimal sets of gene perturbations whose transcriptional responses change the combined state of these pathways from one phenotype to another. Our approach accounts for limited data, heterogeneity within phenotypes, confounding biological variation, and combinatorial explosion in gene sets in ways that traditional methods cannot [13]. In particular, limited data is addressed by our development of the generative model TWAVE; inference of heterogeneous pathways is illustrated in the example of allergic asthma; common drivers of cancer progression across biological subtypes are found in the DepMap example; and finally, a combinatorial explosion is avoided by casting the identification of causal genes as an optimization problem. The approach can also implicate candidate genes through their known downstream effects on the gene regulatory network obtained from experiments.

An overarching goal of our approach is to narrow the scope of candidate gene combinations to a number amenable to targeted low-throughput experiments. As in previous successful applications of Boolean networks [81, 82, 83, 84, 85, 86] and principal component-based techniques that uncover low-dimensional structure in gene regulatory networks [87], we aim to uncover causal influences. The main advantage of our approach is that it can generate predictions solely from publicly available data without explicit network reconstruction or specific knowledge concerning the gene functions and interactions, making it broadly applicable.

It is constructive to reflect on the key assumptions underlying our approach. First, we assume that cellular traits are well reflected by gene expression, which is validated by the fact that we and others [24, 88, 89, 90, 91, 91, 92] can accurately classify gene expression profiles by their phenotypic labels. While transcriptional data do not directly account for post-transcriptional/translational regulation [93], they do account for downstream impacts on gene expression. Nevertheless, it is straightforward to incorporate multi-omic [94, 95, 96, 97] data to directly account for mechanisms beyond transcription. Second, we assume that transcriptional responses combine additively, which has been demonstrated to be a good approximation to control cell behavior [24]. Recent work has applied VAEs to the *forward* problem of estimating nonadditive transcriptional responses to combinatorial perturbations [98], raising the possibility of going beyond the additive assumption in the future. Integration of this technique into our method to solve the *inverse* problem of mapping causes to trait phenotypes, as considered here, would require targeted experiments to train the VAE to recognize nonadditivity. Finally, we assume that our library of transcriptional responses is sufficiently large and diverse to comprehensively capture the impact of genes. Notwithstanding, we demonstrate that enrichment analysis [99, 100] can implicate upstream transcription factors that are not included in our library.

The success of our approach has several far-reaching implications. First, it suggests that cell line perturbations *in vitro* are informative of the gene behavior *in situ* [101]. Second, it shows that the genes needed to drive a phenotypic change are distinct from those that reverse the change. This is a consequence of our optimization model acknowledging the fundamentally different network impacts of reversing a knockdown versus overexpressing a given gene, which is consistent with irreversibility to transient perturbations observed in gene regulatory networks [102, 103]. Third, the success of TWAVE suggests that gene expression can be represented in a low-dimensional space [87], which might be a general feature across many complex network systems [104]. Ultimately, our approach provides a new tool to investigate genotype-phenotype relationships in complex traits, which is applicable across a range of organisms and traits. In humans, our approach also lays a new ground for the design of multi-target disease treatment strategies.

## Materials and Methods

### TWAVE Architecture

We employ a conditional variational autoencoder in PyTorch, which is tailored for the analysis of genomic data and leverages class labels (baseline or variant) to impart enhanced interpretability and classification precision. The architectural blueprint consists of three neural network: an encoder, a decoder, and a classifier.

The encoder comprises two fully connected hidden layers, each embedding 256 and 128 units, respectively, with Rectified Linear Unit (ReLU) activation functions. The input layer takes a single gene expression profile, length aligned with the number of genes, and is concatenated with pertinent class labels. This design not only captures the intricate gene expression patterns but also incorporates class-specific information for a more nuanced latent representation. The decoder component reconstructs the input RNA-Seq data through a series of ReLU-activated layers, culminating in a sigmoid activation function. This reconstruction process aims for the faithful reconstruction of the input profile from the encoded latent space. Simultaneously, the classifier, featuring a linear layer, facilitates class predictions grounded in the extracted latent representation.

### TWAVE Training and Sampling

During the training phase, a set of loss functions drives the optimization process. The combination of reconstruction loss and Kullback-Leibler divergence loss is deployed, ensuring a balance between accurate data reproduction and the regularization of the latent space. The training regimen spans 500−10^4^ epochs, depending on the dataset, with mini-batches consisting of 50−200 samples. We use an Adam optimizer, with a learning rate set at 0.0001.

For sampling, we generate synthetic profiles within the latent space of TWAVE. Latent vectors are obtained by sampling from clusters corresponding to distinct class labels. The latent vectors corresponding to different classes are then clustered, and marginal distributions of baseline and variant profiles in the latent space are extracted using kernel density estimation (KDE). The band-width parameter for our KDE is set to 0.2 to control the smoothness of the estimated density. We then sample new latent space points from these two clusters using our density estimator, and these points are decoded to the full gene expression space with the decoder, producing synthetic profiles for both class labels.

### Bayesian Inference of Causal Eigengenes

We perform Bayesian inference of causal eigengenes from the gene expression data projected onto the top *d* = 200 eigengenes, *X* = *Y U*_*n*×*d*_, where we observe that these eigengenes account for over the overwhelming majority of the variance in all traits. In Fig. S1, we show that the remaining percent variance after *d* = 200 eigengenes is less than 1% for all traits except (lung) cancer metastasis and type 1 diabetes, where the remaining eigengenes account for about 22% and 23.5% of the total variance, respectively. Our choice *d* trims the number of eigengenes to a tractable one that allows for relatively fast Monte-Carlo optimization.

Our inference procedure adapts the fine-mapping of causal variants in GWAS [17] to eigengenes. The first step of the fine-mapping is a logistic regression using class labels. We fit the log-odds ratios *ζ* to the data *X* with effect sizes *β*:

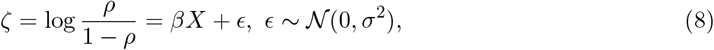

where *ϵ* is a noise vector and *ρ* denotes the probability that the (binary) label is 1. In this scheme, we seek to infer an optimal *d*-dimensional vector of causal effects *γ* = (*γ*_*i*_), *γ*_*i*_ ∈ {0, 1}, which take on values of zero or one depending on whether eigengene *i* is causal or not.

It can be shown that the likelihood of *ζ, X* given *γ* is

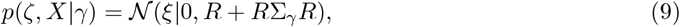

where *ξ* = *X*^*T*^ *ζ/d* is the z-value, 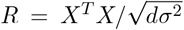 is the eigengene correlation matrix, Σ_*γ*_ = *ds*^2^ diag(*γ*), and *s* is a hyper-parameter. We set an initial *s* = 0.05 as in FINEMAP [17]. Taking a uniform prior on the number of causal effects *k*,

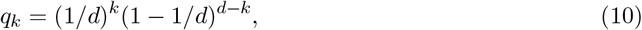

we can then express the posterior distribution of causal effects given our data as

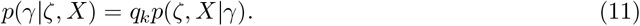

We use MCMC to optimize this distribution over *s* and *γ*, though other techniques such as FINEMAP or the sum of single effects (Susie) model could be employed as well [19]. Posterior inclusion probabilities are calculated as an average over MCMC samples of the posterior distribution

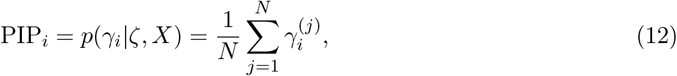

where *N* is the number of samples. The Monte Carlo steps consist of flipping a causal effect (eigengene) on or off at random and we attempt 10^5^ steps with a burn-in period of 10^3^ steps, according to a Metropolis acceptance criterion.

### Maximum Entropy Null Graph Model

As a null model for our gene concurrence graph, we construct a maximum entropy graph constrained by the row and column sums of our matrix *A*. The null model *G* maximizes the entropy *S*(*G*) = − ∑_*G*_ *P* (*G*) ln *P* (*G*), where *P* (*G*) is the canonical distribution *P* (*G*) ∝ exp[−∑_*i*_*β*_*i*_*k*_*i*_ − ∑_*i*_ *γ*_*i*_ *k*_*i*_]. Here, *k*_*i*_ =∑ _*j*_ *A*_*ij*_ is the row sum and *κ*_*i*_ =∑ _*j*_ *A*_*ij*_ is the column sum of node *i*, whereas *β*_*i*_ and *γ*_*i*_ are the respective Lagrange multipliers that enforce *k*_*i*_ and *κ*_*i*_ to be fixed as all other degrees of freedom equilibrate. The row and column sums sequences must follow the maximum entropy conditions

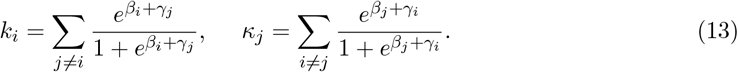

These conditions can be solved for the Lagrange multipliers iteratively as

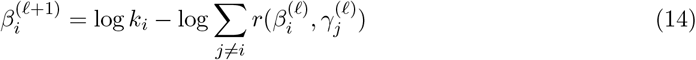

and

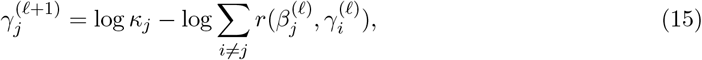

where *l* denotes the iteration and *r*(*x, y*) = 1*/*(*e*^−*y*^ + *e*^*x*^). From here, the link probabilities in Eq. 7 can be computed as

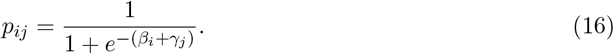

### Complex Disease Data Curation

We obtained our 7 datasets from GEO (see Table 1). The expression matrices, originally in raw counts, are curated keeping those genes and samples that meet the following criterion: average counts in a gene *>* 5 and total counts in a sample *>* 10^5^. The data was normalized to the number of transcripts per million (*N*_TPM_) using reference transcript lengths mapped from gene ENSEMBL id, and the final expression data was saved as log_10_(*N*_TPM_ + 10^−10^) + 10. Labels, whether they are baseline or variant, were one hot encoded.

### Transcriptional Response Library

The transcriptional response data was curated as described in Ref. [24]. The list of GEO series accession numbers and associated gene perturbations are listed in Dataset S1. The inclusion of a knockdown in the library does not imply the inclusion of its overexpression and vice versa.

## Data, Materials, and Software Availability

Raw gene expression counts data are available through GEO. Relevant accession numbers are included in Materials and Methods. The software and processed data for employing the method are available from the GitHub repository: https://github.com/biophysben/Generative-prediction-of-causal-gene-sets-responsible-for-complex-traits. Source data for training TWAVE are stored on Dryad at https://doi.org/10.5061/dryad.s4mw6m9hf.

## Author contributions

B.K.S., T.P.W. and A.E.M. designed research; B.K.S. and B.K.O performed research; B.K.S. analyzed data; and B.K.S., B.K.O., T.P.W., and A.E.M. wrote the paper.

## Author Declaration

The authors declare no competing interests exist.

## Acknowledgments

This work was supported by NIH/NCI grant No. P50-CA221747 through the Malnati Brain Tumor Institute and leveraged research from NSF grant No. MCB-2206974. The authors also acknowledge support from the National Institute for Theory and Mathematics in Biology (NSF Grant No. DMS-2235451 and Simons Foundation Grant No. MP-TMPS-00005320) and the use of Quest High-Performance Computing Cluster at Northwestern University.

## Supporting Information for

### SI Overview

This Supplementary Information file contains a detailed discussion of the choice of the Euclidean metric underlying our optimization, figures that support the analysis presented in the main text, an extended description of the genes associated with changes in trait phenotypes beyond those discussed in the main text, and figures analyzing the results for the pan-cancer, MODY3, and inflammatory bowel disease (IBD) traits. The text contains a detailed discussion of the genes associated with each trait in Table 1 that is not explicitly discussed in the main text, with the corresponding genes laid out in Supplementary Tables S1-S6. Supplementary Figs. S1 and S2 contain numerical and statistical analysis relevant to the identification of causal genes and significant gene pairs, respectively. The results for co-occurring genes in pan-cancer are contained in Supplementary Fig. S3A,B, and the corresponding results for IBD are in Fig. S3C,D. Supplementary Figs. S4-S5 concern a comparison of our method with existing ones in the cases of MODY3 and IBD. The first trait phenotype is purported to be caused by a mutation to a single gene, and in Supplementary Fig. S4 we show that the genes identified by our method overlap with the known downstream effects of that mutation. IBD contains over 1,000 TWAS-associated genes, 44 of which overlap with our perturbations. In Supplementary Fig. S5, we show that the occurrence frequency of a perturbation is associated with the gene being identified by TWAS at a much higher rate than for differentially expressed genes.

#### Justification of the Euclidean metric

In this section, we motivate why we choose to formulate optimization in Eqs. (1)-(2) and (4) of the main text in terms of a Euclidean distance between the baseline and variant states. We recall that the eigengenes are selected for causality via the fine mapping procedure, and thus the Euclidean norm posits that any selected eigengene could equally lead to a change in phenotype. We comment on four representative alternative choices for the distance:

1. Minkowski distances, which allow for *p*—the polynomial order of the norm—to be different from 2 (the Euclidean case);
2. scaled Euclidean metrics, which allow for the different dimensions to be differently weighted;
3. similarity metrics, such as the cosine distance, which characterize the alignment between state vectors; and
4. arbitrary curvilinear metrics, which are well-suited for measuring distance in curved spaces.

Type (1) notions of distance satisfy the triangle inequality when *p ≥* 1, so we focus on those cases. When *p* = 1, the decrease in the objective function is piecewise constant, and the optimal points occur at cusps, but the location of optimal point remains the same. Thus, the perturbations identified by the method presented in the paper is expected to yield similar results, though the value of the regularization parameter *λ* could change. At the same time, the optimization would likely be more expensive due to the large number of non-differentiable points. For *p >* 1, the objective function is smooth, but the optimal point is again the same, so we expect the selected genes to remain largely similar for other values of *p*, although some specifics may change in the process of optimization.

Type (2) notions of distance can readily incorporate the existing data and/or trait-specific biological knowledge to reweight the causal eigengenes. One natural weighting is to rescale the projection along each causal eigengene by the (dataset-wide) standard deviation of expression across that same eigengene. This choice would remove any extensive effects of the selected eigengenes (i.e., eigengenes with more weight among highly-expressed genes would be treated on equal footing with lower-expressed genes). In general, reweighting implicitly reflects a choice that the absolute change in expression of certain genes is more important to the phenotype than others. To remain unbiased as to which absolute changes in gene expression influence phenotype, we did not employ such a reweighting here.

Type (3) notions of distance concern only the direction, but not the magnitude of the changes. However, as far as cell behavior is concerned, both the magnitude and direction of the change are expected to be important. We explicitly account for this by restricting the range of perturbation strengths we allow in Eqs. (1)-(2) and (4) of the main text, and we find cases in which multiple perturbations that point in almost the same direction in transcriptional space are needed to realize a change in the cell behavior. Type (4) notions of distance require the specification of a global metric tensor at all points in transcriptional space, which implicitly makes substantial assumptions about the transcriptional regulatory network that are difficult to verify with the existing data. As a result, we eschewed implementing metrics of this type in Eqs. (1)-(2) and (4) of the main text.

Speaking loosely, the Euclidean metric is a parsimonious choice that treats the causal eigengenes on equal footing.

#### Genes implicated in complex traits

We now discuss the remaining complex traits mentioned in Table 1 but not otherwise discussed in detail in the main text, and the gene perturbations our method associates with their phenotypes. We recall that we use italicized capital letters to refer to genes, while unitalicized capital letters are used for acronyms denoting traits.

##### Inflammatory bowel disease

IBD, which includes ulcerative colitis and Crohn’s disease, is a chronic intestinal inflammatory disease associated with immune system dysregulation in which an imbalance occurs between anti-inflammatory and pro-inflammatory responses (S1). Accordingly, some of the perturbations that account for the largest transcriptional differences between the average healthy transcriptional profile and the average IBD-associated transcriptional profile are associated with genes involved in immune system function (see Table S1).

Several of the genes in Table S1 have also been found in the literature to be differentially expressed in inflammatory tissues. These genes include:

1. *CPSF3*, which has been found to be markedly higher in ulcerative colitis tissues than in healthy tissues (S1);
2. *CD86*, which is upregulated in B-cells of mouse models and human patients with IBD (S2);
3. *PROX1*, which is upregulated in Crohn’s disease patients (S3); and
4. *CDK12*, which is often downregulated to mitigate inflammatory disease (S4).

All of these genes make sense biologically since they are known to have roles in regulating immune function.

#### Food allergy

Food allergies are a sustained overreaction of the immune system to harmless environmental factors, with disruptions in early immune development likely playing a role (S5; S6). Our method implicates several genes crucial to immune system development and immune response modulation, including *CEBPA, LEF1, ERG, TCF7L1, ONECUT2, ETV1*, and *CBFB* (see Table S2). Further, *LEF1* coexpression with *TCF1* (which is in the same protein family as *TCF7L1* ) has been shown to lead to chronic activation of helper T-cells, likely due to their combined roles in the Wnt pathway critical for T-cell memory (S5).

#### Cancer metastasis

Cancer cells undergo a series of transcriptomic and physiological changes that give rise to metastases in different tissues, with the cells’ phenotypic plasticity playing a role (S7). Consistent with this observation, we find that genes related to countervailing processes of the epithelial to mesenchymal transition (EMT) and tumor suppression are implicated in cancer metastasis by our method (Table S3). For example, our method identifies *PTHLH*, which is associated with cell migration (S8). In other cases, our method identifies related genes whose polarity of perturbation (up or down regulation) runs counter to that observed in previous experiments. Such genes include *FAM83H-AS1*, also associated with cell migration (S9), and *MYCN*, which is associated with tumor proliferation (S10). In these instances, the same cellular processes are targeted, but outside factors may alter the way that the specific gene upregulations and downregulations influence phenotype.

#### Age-related macular degeneration

Age-related macular degeneration (AMD) is a leading cause of vision impairment and loss (S11). Disease progression is characterized by the build-up of extracellular deposits in the macula (an anatomical structure within the retina of the eye) and the ensuing degeneration of photoreceptors and nearby tissue (S12). Dysregulation of lipid, vascular, inflammatory, and extracellular matrix pathways have been implicated in this disease (S11). Accordingly, several of the perturbations for AMD identified in Table S4 are in genes involved in these pathways. Specifically, *MIR29A* was found to disrupt the formation of blood vessels in the eye (S13), while a polymorphism in *IGF1R* (a gene with a role in inflammatory responses and angiogenesis) is significantly associated with advanced AMD (S14).

#### Type 1 diabetes

Type 1 diabetes (T1D) progresses through autoimmune attacks on insulin-producing *β*-cells within the pancreas (S15). In agreement with this, our method identifies perturbations to genes involved in immune development and response, like *MIR126, CEBPA*, and *ZAP70*, as causal in T1D. Interestingly, we also find perturbations to genes with functions relevant to the pathogenesis of type 2 diabetes (T2D), including insulin resistance and storage and metabolic breakdown of fats (S16). We further find perturbations to genes involved in vascularization, as in processes known to be disrupted as part of the long-term complications of T2D (S16).

#### Non-small cell lung cancer

Non-small cell lung cancer (NSCLC) represents 85% of new lung cancer cases (S17). Our predicted perturbations involve several genes associated with cell growth, proliferation, and tumor suppression. These include *EGFR*, which is expressed in a variety of human tumors, including NSCLC (S18). Furthermore, *BCL11B* has been shown to have tumor suppressing functionality (S19).

## Supplementary Figures

**Figure S1:**
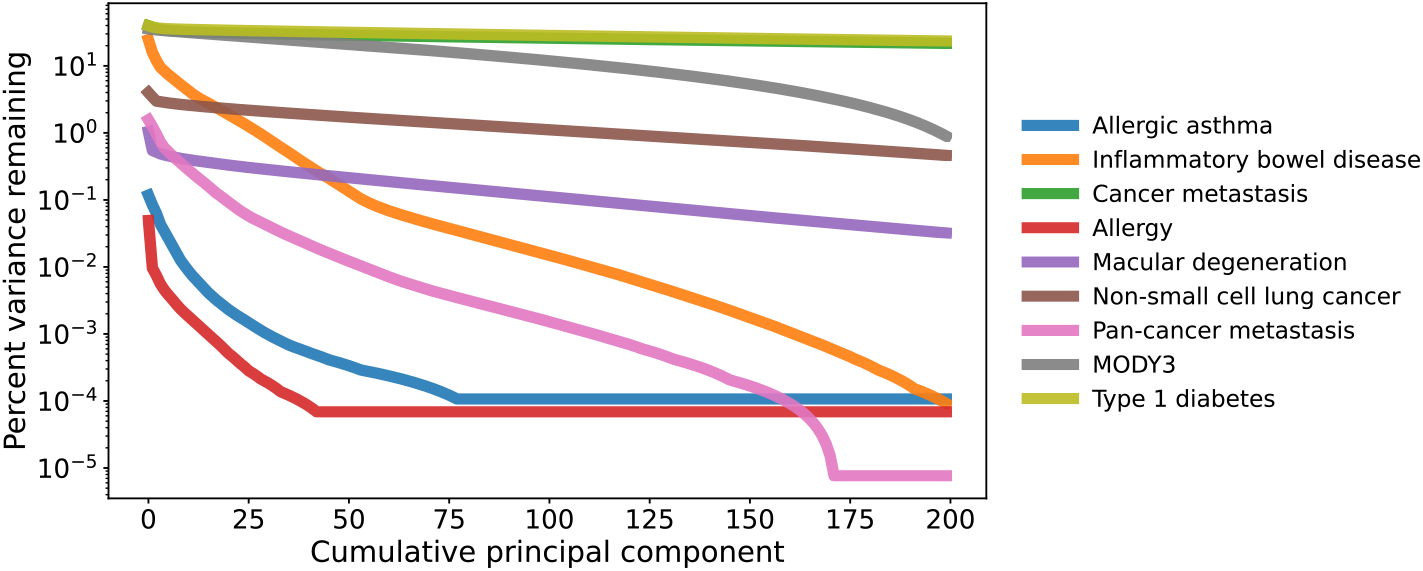
Justification for using *d* = 200 eigengenes. The percent of unexplained variance in the data is plotted as a function of the number of principal components (ordered by the fraction of variance explained). The top 200 eigengenes explain *>* 99% variance for all traits, except cancer metastasis and type 1 diabetes, where *>* 77% of the variance is explained.

**Figure S2:**
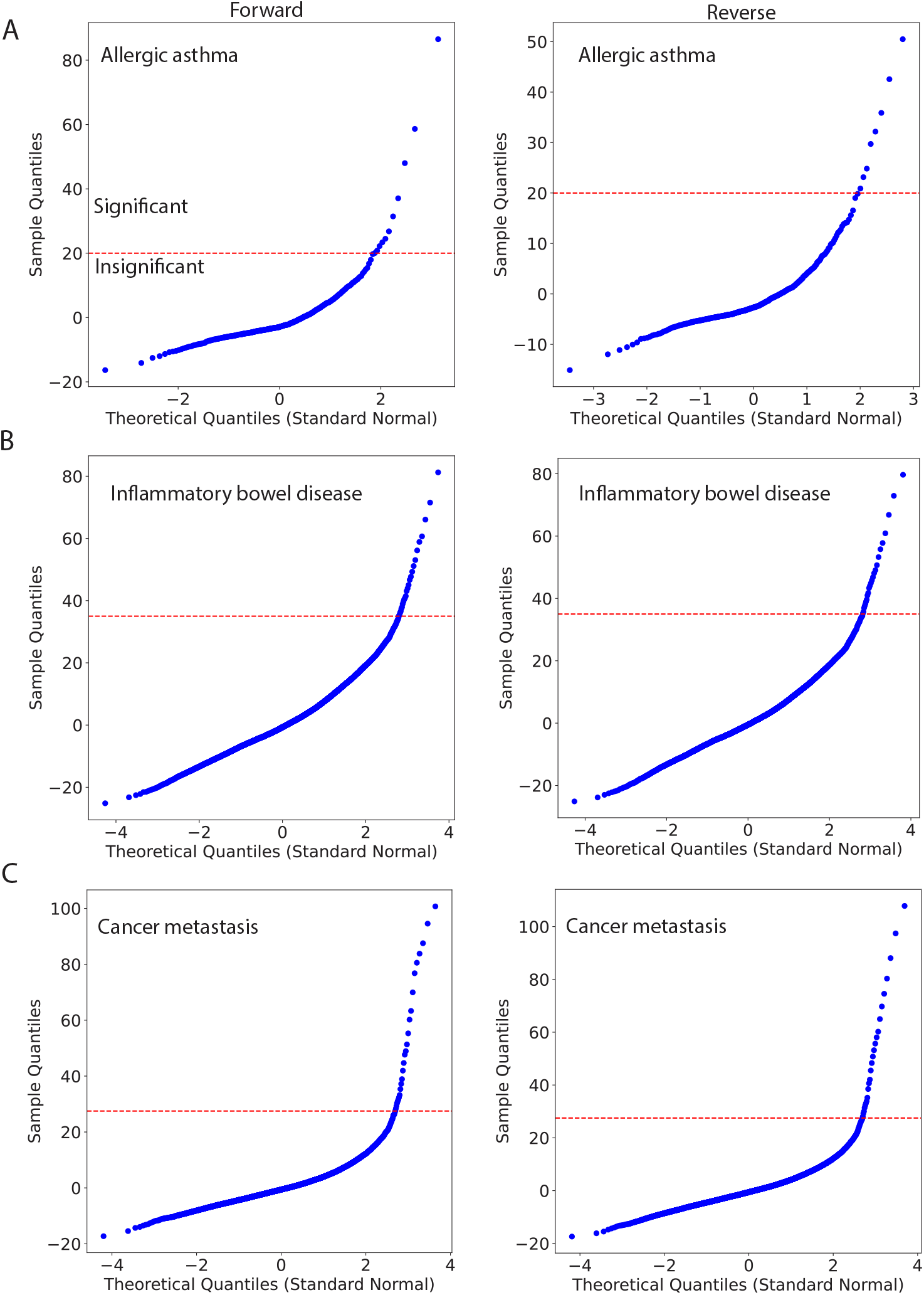
Quantile-quantile plots for allergic asthma (*A*), inflammatory bowel disease (*B* ) and pan-cancer metastasis (*C* ). Horizontal lines indicate thresholds above which co-occurring gene pairs are considered significant. Each case shows the baseline-to-variant (left) and reverse (right) directions.

**Figure S3:**
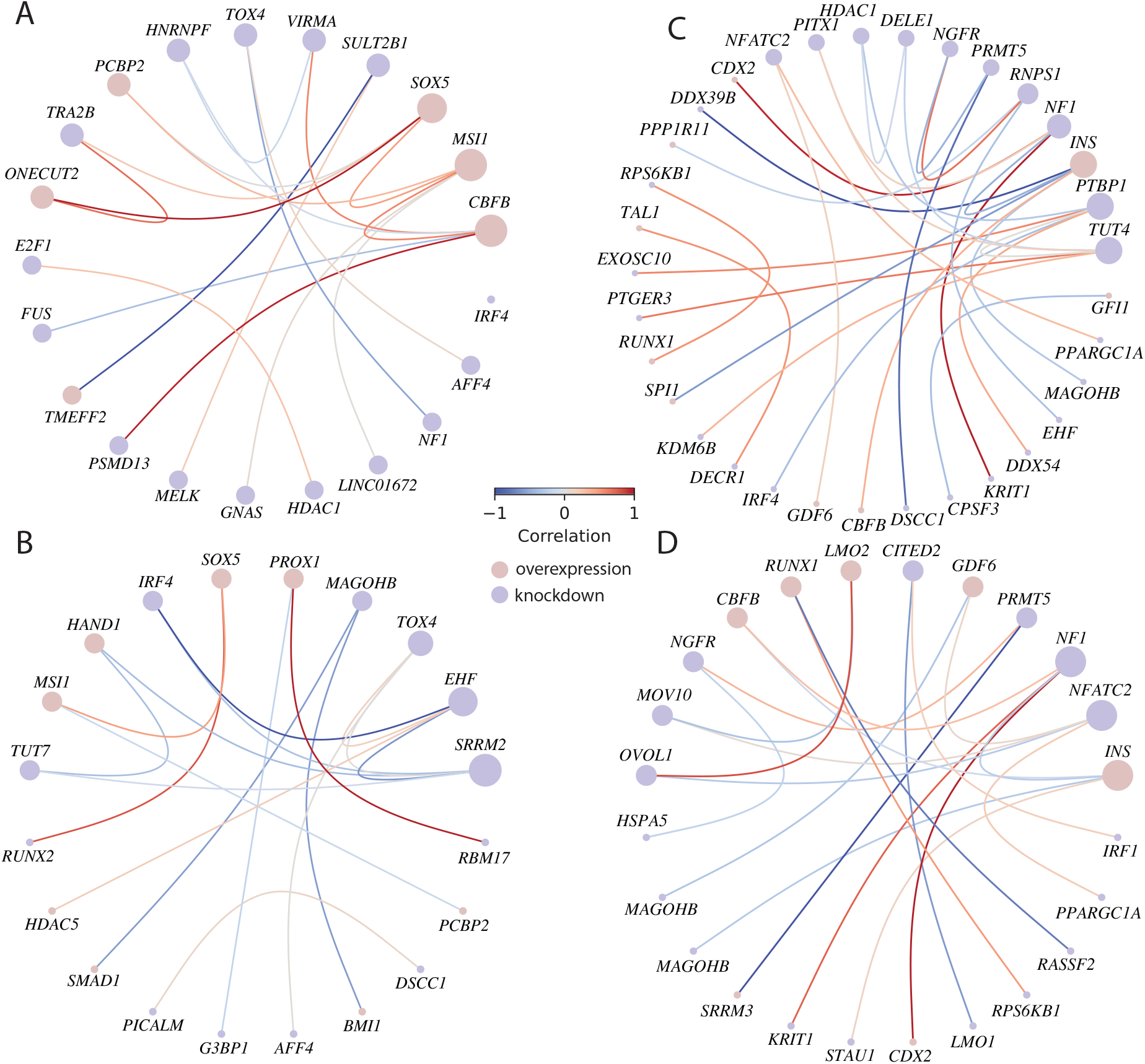
Gene pair co-occurrence networks for (*A, B* ) pan-cancer metastasis and (*C, D* ) inflammatory bowel disease. The baseline-to-variant pairs are indicated in (A, C) and reverse pairs in (B, D). The edge colors indicate correlations between the genes, the node colors indicate the type of perturbation, and the node sizes are proportional to the number of edges.

**Figure S4:**
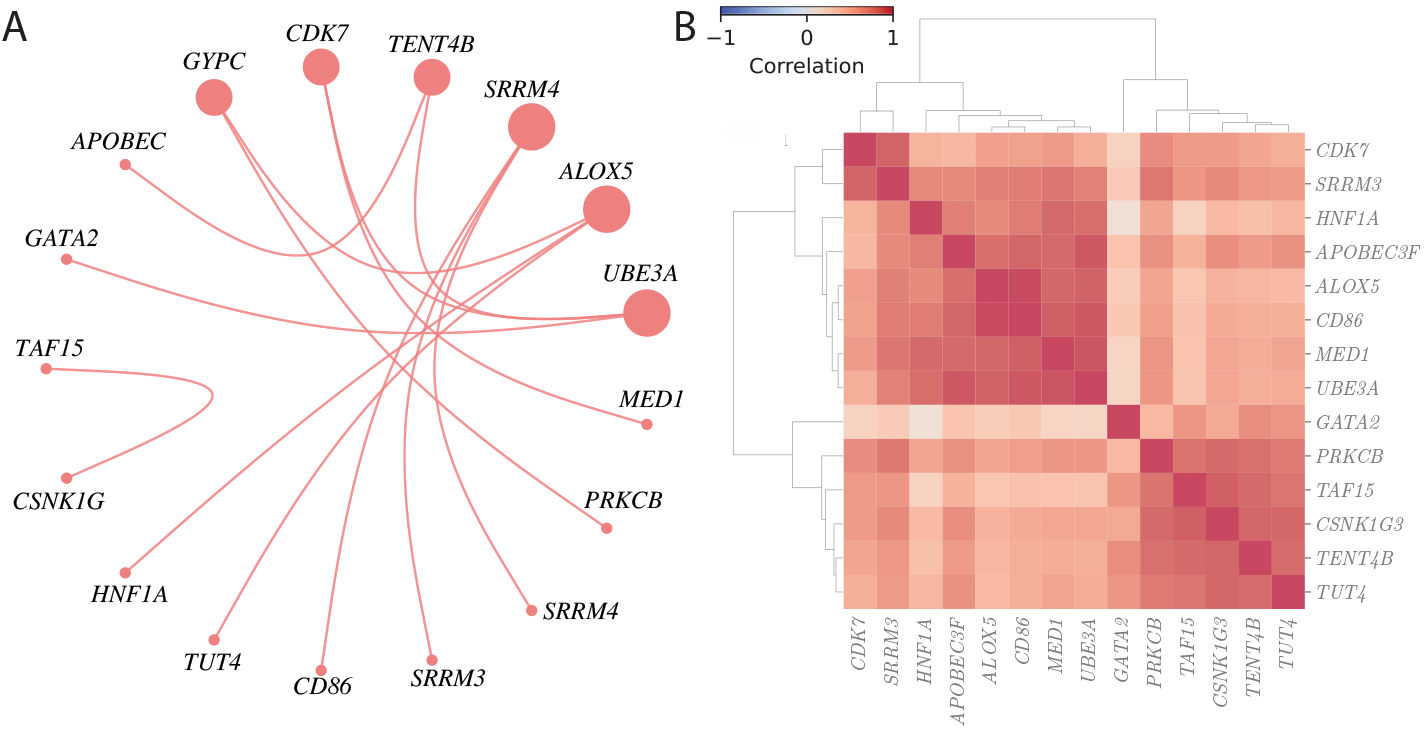
Co-occurring genes in the MODY3 trait and the correlations between them. (*A*) Gene co-occurrence network in the baseline-to-variant direction. (*B* ) Color-coded gene expression correlations of the genes in (A) across all transcriptional responses in the library. The transcriptional responses to *SRRM4* and *GYPC* appear in the library but their expressions are not measured in the dataset, so they do not appear in (B).

**Figure S5:**
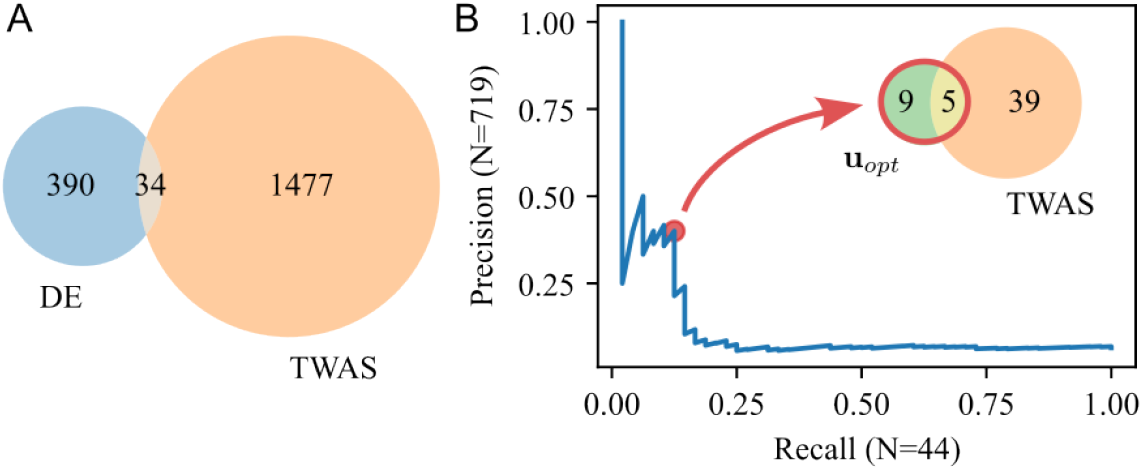
Comparison of genes identified by differential expression, *u*_*opt*_ (our method), and TWAS for inflammatory bowel disease. (*A*) Venn diagram of the differentially expressed genes in GSE193677 (S20) and all TWAS genes associated with inflammatory bowel disease. Only 8% of the differentially expressed (DE) genes are also implicated by TWAS. (*B* ) Precision-recall curve of the 719 unique genes associated with columns of the perturbation matrix *B* versus the 44 TWAS genes that overlap with these perturbed genes. Genes were ranked by the number of times they appear in the point-to-point forward optimizations for IBD. For the top *n* genes, the precision is the fraction of the *n* genes that are in the set of TWAS genes, while the recall is the fraction of the 44 TWAS genes contained in the top *n* genes. The red dot indicates *n* = 14 genes of which 5 overlap with TWAS, as indicated in the Venn diagram.

## Supplementary Tables

**Table S1:** Transcriptional perturbations associated with inflammatory bowel disease.

| Gene | Annotation |
| --- | --- |
| *CPSF3*<sup>-</sup> | post-transcriptional processing (S21) |
| *UBE2O*<sup>-</sup> | protein quality control, blood cell differentiation (S22) |
| *CD86*<sup>-</sup> | T-cell activation (S23) |
| *PROX1*<sup>+</sup> | developmental transcription factor (S3) |
| *CDK12*<sup>-</sup> | cell cycle progression (S24) |
| *ADARB1*<sup>-</sup> | adenosine-to-inosine (A-to-I) RNA editing (S25) |
| *RBJ*<sup>-</sup> | GTPase signaling protein (S26) |
| *SLIRP*<sup>-</sup> | RNA binding protein (S27) |
| *NOTCH1*<sup>+</sup> | intracellular signaling, cell fate (S28) |
| *SOX2*<sup>-</sup> | embryonic development (S29) |
| *RNF43*<sup>+</sup> | DnaJ heat shock, regulation of Wnt signaling, tumor suppression protein (S30) |
| *NAB2*<sup>+</sup> | transcriptional corepressor of EGR genes (S31) |

**Table S2:** Transcriptional perturbations associated with food allergy.

| Gene | Annotation |
| --- | --- |
| *CEBPA*<sup>+</sup> | transcription factor, roles in adipogenesis, immune cell differentiation, metabolic homeostasis (S32; S33) |
| *LINC00673*<sup>-</sup> | expression regulation, tumor suppression, involved in cancer progression (S34) |
| *LEF1*<sup>+</sup> | transcription factor in the Wnt/β-catenin signaling pathway, hematopoiesis, immune response modulation (S35; S5) |
| *ERG*<sup>+</sup> | transcription factor in the ETS family, with roles in immune cell differentiation (S36) |
| *ADARB1*<sup>-</sup> | adenosine-to-inosine (A-to-I) RNA editing (S25) |
| *TCF7L1*<sup>-</sup> | regulator in the Wnt/β-catenin signaling pathway—roles in T-cell differentiation, epithelial barrier function, and immune response (S37) |
| *ONECUT2*<sup>-</sup> | epithelial differentiation, immune cell regulation, and tissue development (S38) |
| *RBPJ*<sup>-</sup> | transcriptional mediator of Notch signaling—differentiation, inflammation, and tissue homeostasis (S39) |
| *ETV1*<sup>-</sup> | transcription factor from ETS family, known for role in cell differentiation, immune regulation, and tissue homeostasis (S40) |
| *MSI1*<sup>+</sup> | RNA-binding protein, with roles in stem cell maintenance, epithelial regeneration, and immune cell function (S41) |
| *CBFB*<sup>-</sup> | regulates immune cell development, osteogenesis, hematopoiesis (S42) |
| *BMPR2*<sup>-</sup> | receptor in the TGF-β/BMP signaling pathway, which regulates immune responses, tissue homeostasis, and inflammation (S43) |

**Table S3:**
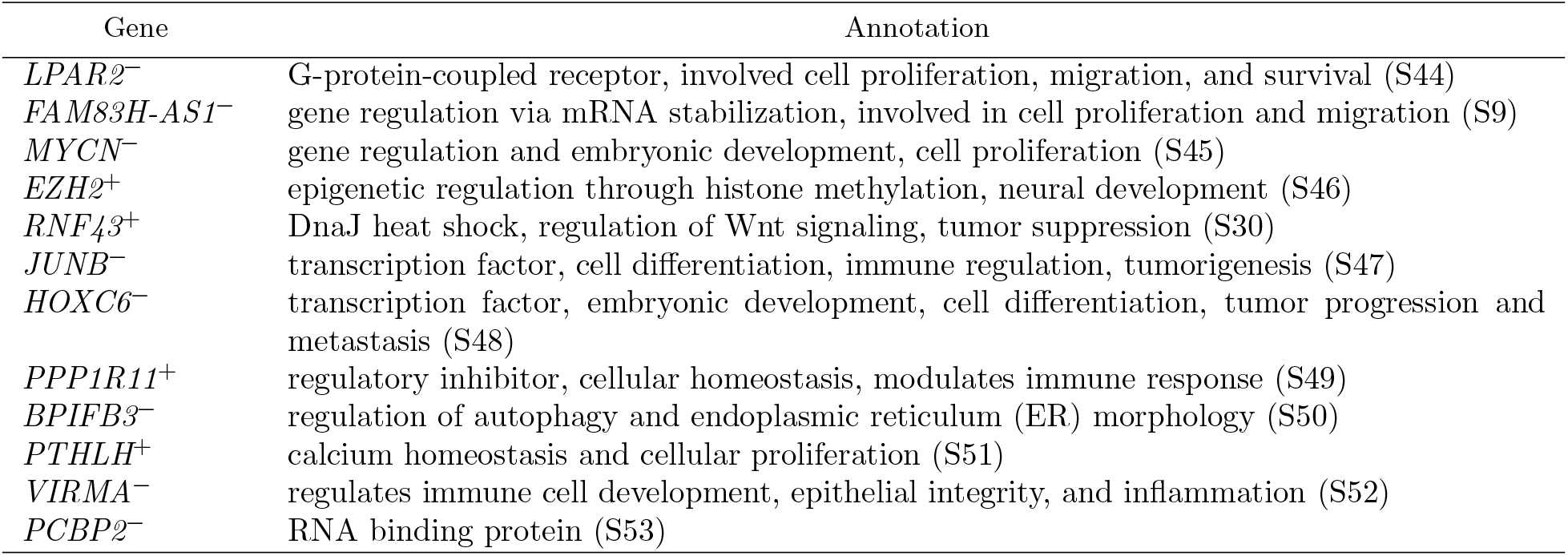
Transcriptional perturbations associated with cancer metastasis to the lung.

**Table S4:**
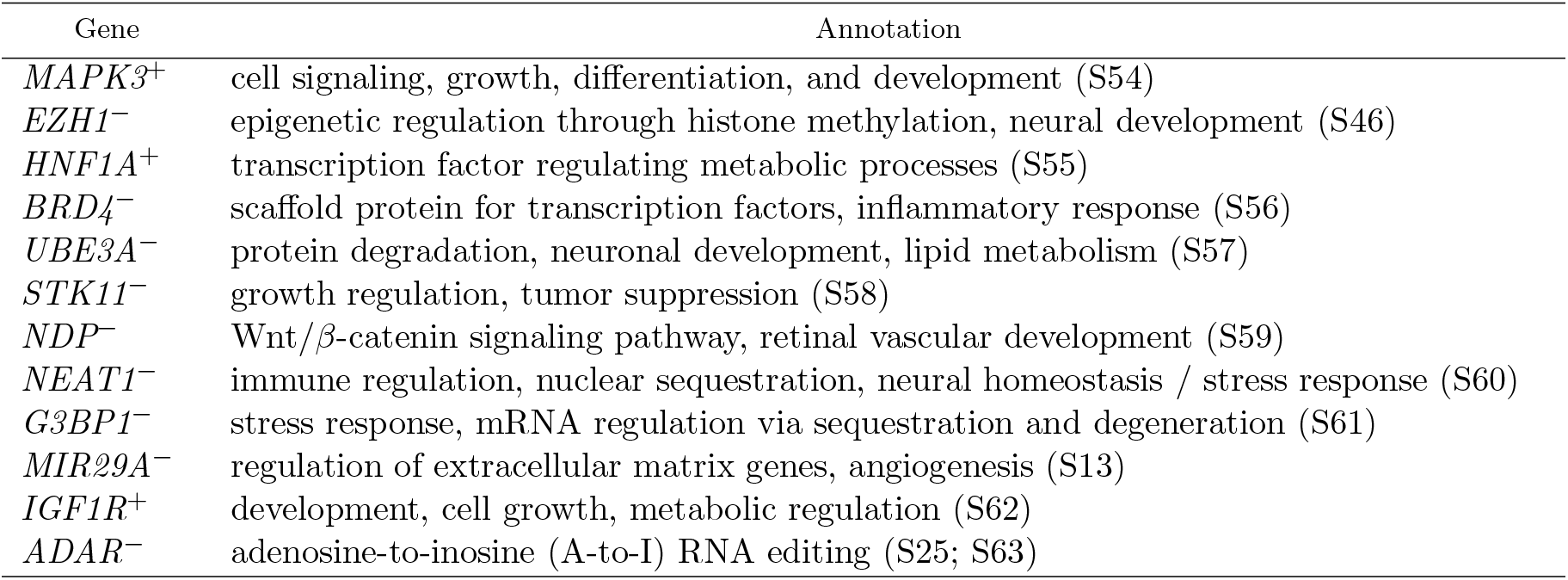
Transcriptional perturbations associated with age-related macular degeneration.

| Gene | Annotation |
| --- | --- |
| *MAPK3*<sup>+</sup> | cell signaling, growth, differentiation, and development (S54) |
| *EZH1*<sup>-</sup> | epigenetic regulation through histone methylation, neural development (S46) |
| *HNF1A*<sup>+</sup> | transcription factor regulating metabolic processes (S55) |
| *BRD4*<sup>-</sup> | scaffold protein for transcription factors, inflammatory response (S56) |
| *UBE3A*<sup>-</sup> | protein degradation, neuronal development, lipid metabolism (S57) |
| *STK11*<sup>-</sup> | growth regulation, tumor suppression (S58) |
| *NDP*<sup>-</sup> | Wnt/β-catenin signaling pathway, retinal vascular development (S59) |
| *NEAT1*<sup>-</sup> | immune regulation, nuclear sequestration, neural homeostasis / stress response (S60) |
| *G3BP1*<sup>-</sup> | stress response, mRNA regulation via sequestration and degeneration (S61) |
| *MIR29A*<sup>-</sup> | regulation of extracellular matrix genes, angiogenesis (S13) |
| *IGF1R*<sup>+</sup> | development, cell growth, metabolic regulation (S62) |
| *ADAR*<sup>-</sup> | adenosine-to-inosine (A-to-I) RNA editing (S25; S63) |

**Table S5:** Transcriptional perturbations associated with type 1 diabetes.

| Gene | Annotation |
| --- | --- |
| *COP1*<sup>-</sup> | insulin secretion, protein degradation, tumor suppression (S64) |
| *UBE3A*<sup>-</sup> | protein degradation, neuronal development, lipid metabolism (S57) |
| *PHF8*<sup>+</sup> | epigenetic regulation, neuronal development (S65) |
| *RNF20*<sup>-</sup> | epigenetic regulation, adipogenesis (S66) |
| *MIR126*<sup>+</sup> | epigenetic regulation of metabolism, inflammatory response, angiogenesis (S67) |
| *CEBPA*<sup>+</sup> | transcription factor, roles in adipogenesis, immune cell differentiation, metabolic homeostasis (S32; S33) |
| *C9orf72*<sup>-</sup> | metabolic homeostasis, motor neuron function (S68) |
| *BMPR2*<sup>-</sup> | osteogenesis, cell differentiation, vascular function (S43) |
| *MOV10*<sup>-</sup> | gene regulation, development (S69) |
| *MIR29B1*<sup>-</sup> | regulation of extracellular matrix genes, promotion of endothelial function (S70; S71) |
| *ZAP70*<sup>-</sup> | immune regulation and development (S72) |
| *BMI1*<sup>+</sup> | transcriptional repressor, mediation of signaling pathways in cardiovascular tissues (S73) |

**Table S6:**
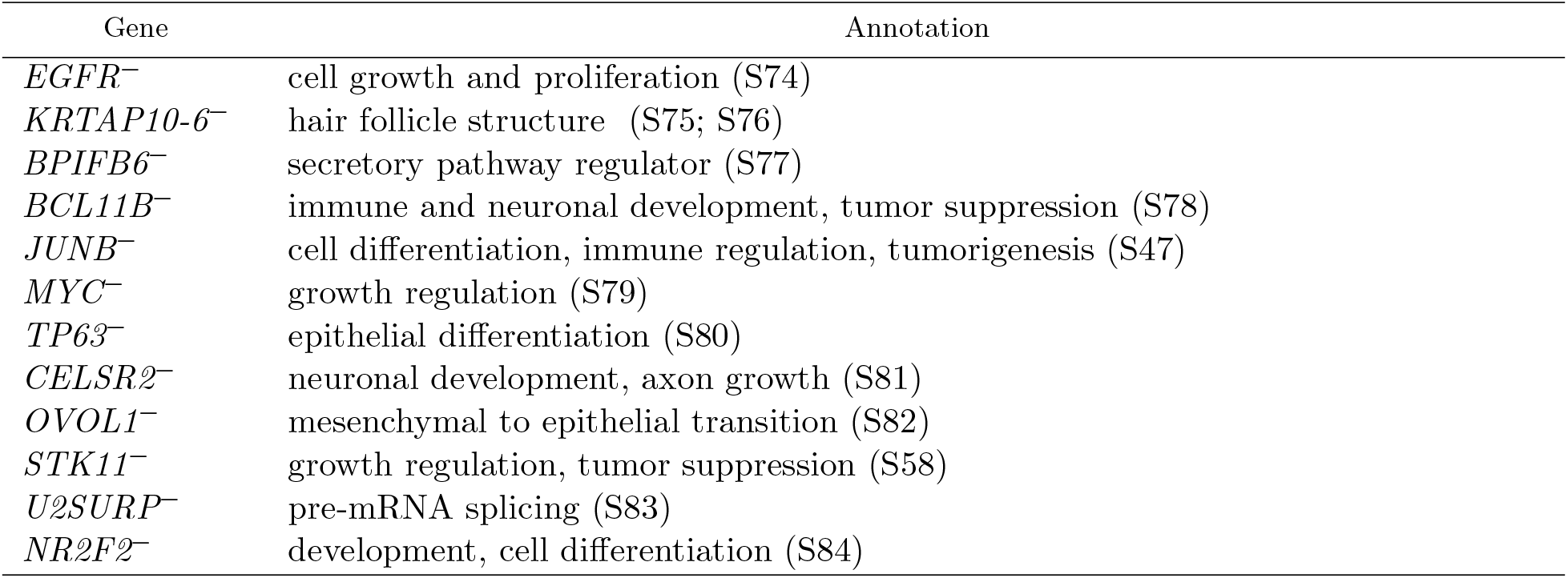
Transcriptional perturbations associated with non-small cell lung cancer.

## Supplementary Datasets

### Dataset_S1-transcriptional_response_metadata.xlsx

Excel file of metadata including the SRA accession numbers of the publicly available data used to specify the transcriptional response matrix *B*. The NCBI Bioproject, cell line, cell type, treatment, and genotype of the initial (unperturbed) and treated (perturbed) states are recorded (left to right) in this table.

